# Defining mononuclear phagocyte distribution and behaviour in the zebrafish heart

**DOI:** 10.1101/2024.04.12.589185

**Authors:** Bethany Moyse, Joanna Moss, Laura Bevan, Aaron Scott, Valérie Wittamer, Rebecca J. Richardson

## Abstract

Mononuclear phagocytes (MNPs) are recognised as highly plastic, multifunctional cells that influence multiple physiological and pathophysiological states. In the heart, they support homeostatic functions, contribute to disease progression and play multiple roles in reparative and regenerative processes following tissue damage. Understanding the heterogeneous populations of cells that contribute to these diverse functions is crucial to facilitating beneficial, and limiting adverse, cardiac outcomes. However, characterisation of precise populations of cardiac immune cells remains incomplete in vertebrate models capable of endogenous regeneration, such as adult zebrafish. Here, we use a combination of transgenic lines to identify distinct MNPs in the zebrafish heart. We show that larval macrophage populations have different origins and a sub-population of *csf1ra* expressing cells are maintained on the surface of the adult heart. MNPs are differently distributed in the myocardium, exhibit different behaviours and are distinguished via expression level of *csf1ra* and *mpeg1.1*. Following injury, tissue resident macrophages rapidly proliferate potentially contributing to reduced scarring. The adult zebrafish heart contains multiple populations of MNPs that can be defined by existing tools. This new understanding of innate immune cell populations in the heart of adult zebrafish sheds light on the composition of a pro-regenerative cardiac microenvironment.

## Introduction

Cells of the mononuclear phagocyte system, namely monocytes, macrophages and dendritic cells (DCs), can be defined by their tissue distribution, function and morphology. Mammalian classical monocytes are short-lived cells that are recruited to sites of inflammation via the vasculature where they differentiate into monocyte-derived macrophages (MDMs) or monocyte-derived DCs^1^. Macrophages are found exclusively within tissues and are derived from adult monocyte populations or self-renewing embryonic-derived tissue resident populations^1^. These cells have an abundance of effector functions, including phagocytosis, macropinocytosis, antigen presentation and the secretion of inflammatory and trophic factors^2^. DCs have unique and primary functions in immunomodulation, processing and presenting antigens to adaptive immune cells within lymphoid tissues, serving to coordinate the innate and adaptive arms of the immune system^2,3^. DCs share many functional, morphological and transcriptional characteristics with macrophages making separation of these cells complex^4^.

In addition to their recruitment to sites of inflammation via monocyte precursors, extensive populations of macrophages, and smaller populations of DCs, exist within tissues during the steady state^5^. Here, the plasticity of macrophages allows them to adopt unique, tissue specific functions. For example, cardiac macrophages possess important roles in supporting the electrical conductance of the heart, whilst microglia promote maintenance of healthy neural networks^6,7^. It was previously thought that tissue resident macrophages were constantly replenished by circulating monocytes. However, it has become evident that in most mammalian tissues, macrophage origin is both diverse and tissue dependent, and consequently influences cellular function^1,8,9^.

Vertebrates exhibit multiple, conserved waves of haematopoiesis^10–12^. During the initial, primitive wave, macrophages emerge from a monopotent progenitor and migrate into tissues to seed initial tissue macrophage populations^10,13^. In mice, primitive haematopoiesis occurs in the yolk-sac blood islands and, correspondingly, initiates in the anterior lateral mesoderm (ALM) in zebrafish^10,13,14^. Primitive haematopoiesis is succeeded by an interim phase when MNPs derive from a transient erythromyeloid progenitor (EMP). Finally, haematopoietic stem cells (HSCs) arise during definitive haematopoiesis and continue as the sole source of monocyte-derived MNP production throughout the lifespan of the organism^1,10,11^.

The homeostatic murine heart has extensive populations of cardiac tissue macrophages (cTMs) dispersed within the myocardium and surrounding endothelial cells^15–18^. Lineage tracing of murine cTMs has shown that primitive macrophages initially colonise the myocardium, followed by definitive, foetal-liver-derived macrophages. Both populations proliferate to maintain cTMs throughout the lifespan of the mouse, with little contribution from circulating monocytes^15,17,18^. These cTMs have an M2-like profile and function to phagocytose cellular debris, maintain the extensive vasculature and modulate the inflammatory environment^15,19^. Following myocardial infarction, these embryonic lineages have distinct responses to tissue injury compared to MDMs, and their dynamics differ between regenerative neonatal and non-regenerative adult mice. Embryonic cTMs are notably distinguished from MDMs by the absence of the cell surface receptor, CCR2^9,18,19^. In neonatal mice, CCR2-embryonic cTMs expand and mediate tissue regeneration, and there is minimal recruitment of CCR2+ monocytes^20^. In contrast, cardiac injury in adult mice induces a profound recruitment of CCR2+ monocytes, which infiltrate the myocardium and differentiate into CCR2+ MDMs to replace embryonic cTMs. These CCR2+ MNPs drive harmful pro-fibrotic activity in contrast to embryonic populations^20,21^ and this is partly attributed to the opposing inflammatory profile of embryonic cTMs and CCR2+ MNPs. Embryonic cTMs retain a principally M2-like expression profile whereas CCR2+ monocytes show elevated expression of pro-inflammatory markers, including Il-1β, Ccl2 and Il6^9,20,21^. Similarly, the retention of pro-inflammatory CCR2+ MNPs has been repeatedly shown to drive adverse cardiac remodelling in the mouse^9,22,23^. Manipulating CCR2+ monocytes may therefore serve as a therapeutic strategy to promote cardiac regeneration in adult mammals.

Similarly, we and others have shown that regenerative zebrafish also exhibit a robust inflammatory response to tissue injury, including rapid upregulation of *tnfa* and *il-1β*^24–26^. However, multiple studies have shown that macrophages are crucial for the regenerative capacity observed in zebrafish tissues^24–31^ and our previous work has started to reveal specific roles for different sub-populations^24^. A wealth of species-specific cell surface and transcriptomic markers have been established to stratify mammalian monocytes, macrophages, DCs, and sub-populations within these lineages^2^. On the contrary, although analogous populations have been described in zebrafish^32,33^, there are currently limited tools available to delineate MNP populations^34,35^ and our understanding of their different, and potentially pro-regenerative, functions remains limited.

Nevertheless, by exploiting the genetic tractability and imaging capabilities of the transparent larval zebrafish, transgenic lines have greatly facilitated our understanding of the *in vivo* dynamics of macrophages. Established zebrafish macrophage markers include *csf1ra*^36^, *cxcr3.2*, *marco, ptpn6*^37^, *mfap4*^38^ and *mpeg1.1* (formerly known as *mpeg1*)^39^. However, much of the initial characterisation of these markers was performed in larval zebrafish and, although expression in tissue macrophages was confirmed, whether expression extends to monocytes and DCs has not been well defined. Recent technological advancements and accessibility to RNA sequencing technology has facilitated the molecular characterisation of MNPs^40–43^. However, this analysis is in its infancy, and few of the uncovered markers have been translated into tools which can be utilised to perform functional analysis of cells *in vivo*. Given the vast heterogeneity of MNPs, there is therefore still much to discover about how to segregate such populations in zebrafish, which will undoubtedly help to reveal the mechanisms by which precise MNPs facilitate regeneration.

## Methods

### Zebrafish lines and maintenance

All zebrafish lines were maintained, and procedures performed, in accordance with guidelines from the UK Home Office and the Directive 2010/63/EU of the European Parliament on the protection of animals used for scientific purposes. All experiments conformed to local University regulations. Adult zebrafish, aged 6-18 months, were randomly selected for analysis/experimental procedures from mixed-sex tanks housing up to 20 individuals. Animals were euthanised using the Schedule 1 method of immersion in an overdose of MS-222 anaesthetic in aquarium water. The TgBAC(*csf1ra:GFP*)^44^, Tg(*mpeg1.1:mCherry*)^39^, Tg(*mpeg1.1:eGFP*)^gl22^[^39^], Tg(*myl7:GFP*)^45^, Tg(*myl7:HRAS-mCherry*)^46^, TgBAC(*csf1ra:Gal4i)*^47^; Tg(*UAS:kaede*)*s1999t*^48^*, csf1ra^j4e1/j4e1^* [^36^], *cmyb^t25127^* [^49^], Tg(*kdrl:mCherry-CAAX*)^50^ and Tg(*kdrl:cre*)^s898^[^51^], Tg(*actb2:loxP-STOP-loxP-DsR^edexpress^*)sd5[^51^] fish have been described previously.

Cardiac injuries on adult zebrafish were carried out as described previously^52^. Briefly, animals were anaesthetised via immersion in 0.025% MS-222 and a ∼4 mm incision made through the skin and the pericardial sac directly above the heart using a microscalpel. The exposed ventricle was dried using a sterile cotton swab and a liquid nitrogen cooled metal probe was applied for 30 seconds. Fish were returned to aquarium water and allowed to fully recover before being transferred back to the aquarium until the desired timepoint.

### Wholemount immunostaining on larvae

Larvae and juveniles at 2-24 days post fertilisation (dpf) were fixed in 4% paraformaldehyde (PFA) (Sigma; P6148) overnight at 4°C, washed in PBS and permeabilised in 0.25% Trypsin for 15 minutes on ice. Larvae were washed three times in PBS-Tx (0.02% Triton-X in PBS), transferred to blocking buffer (5% horse serum in PBS) for 2 hours and incubated in primary antibody (anti-Myosin heavy chain, A4.1025 DHSB, 1:250; anti-L-plastin^53^, 1:500) overnight at 4°C. Samples were washed in PBS-Tx and blocked for 1 hour in blocking buffer and stained with secondary antibody (Alexa Fluo-647, SA5-35500 Thermo Fisher Scientific, 1:500) for 2 hours. Samples were mounted laterally in 1% agarose.

For whole mount imaging of adult hearts, tissues were dissected and washed in ice-cold PBS (ThermoFisher; 18912014; PBS) supplemented with 10 U/ml heparin (Alfa Aesar; A16198) and fixed in 4% PFA at 4°C overnight or at room temperature for 2 hours on a rotator. Tissues were further washed in PBS prior to mounting in 1% low gelling temperature agarose and imaging.

### Immunostaining and Ce3D tissue clearing of adult hearts

Ce3D tissue clearing was performed using a previously established method^54^. Tissues were protected from light at all stages. Tissues were prepared, fixed, and immunostained prior to clearing. Briefly, fixed hearts were washed in conventional washing buffer^54^ and incubated in conventional blocking buffer^54^ on a shaker at 150–220 rpm at 37 °C for 8–24 hours or at room temperature for 48 hours. Tissues were subsequently incubated with antibodies (anti-Phospho-Histone H3 (Ser10), Cell Signaling Technology-3377; anti-GFP, Abcam-ab13970; anti-mCherry, Invitrogen-M11217) diluted in conventional blocking buffer on a shaker at 150–220 rpm for 3-4 days at 37 °C. Tissues were washed in conventional washing buffer at 37 °C for 8–14 h then overnight at room temperature. To clear tissues, hearts were dried to eliminate washing buffer then incubated in Ce3D clearing solution^54^ on a rotator at room temperature until clear (∼1-2 days). For imaging, hearts were mounted in Ce3D clearing solution in 35 mm glass bottom CELLview culture dishes (Greiner; 627861).

### Imaging

Imaging of whole mount larvae and adult hearts was performed on a Leica TCS SP8 AOBS confocal laser scanning microscope using a 10x/0.4 HC PL APO Dry or 20x/0.75 HC PL APO CS2 Immersion objective.

For *ex vivo* imaging of excised hearts a previously published protocol^55^ was adapted. Adult Tg(*mpeg1.1:mCherry*); TgBAC(*csf1ra:GFP*) zebrafish were euthanised and the heart extracted (preserving the ventricle, bulbus arteriosus and atrium) into 0.22 µm filter-sterilised and pre-warmed (28°C) culture media (Dulbecco’s Modified Eagle’s Medium – High Glucose (Sigma; D5796-500ml), 10% Cytiva HyClone™ Fetal Bovine Serum (Fisher Scientific; 11591821), 1% MEM Non-Essential Amino Acids Solution (ThermoFisher Scientific; 11140050), 100 U/ml Penicillin-Streptomycin, 50 μM 2-mercaptoethanol (Fisher; 31350010) and 100 μg/ml Normocin (InvivoGen; ant-nr-1)). 30 mM 2,3-butanedione monoxime (BDM) was omitted as it has been shown to inhibit leukocyte migration^56^. Within a laminar flow hood, hearts were transferred to a 1.5 ml Eppendorf tube using a P1000 with a large aperture tip and washed three times with PBS. For imaging, a circular mould was made inside a 35 mm glass bottom CELLview culture dish (Greiner; 627861) and one drop of 1% molten agarose added in the base. Hearts were transferred on top of this before mounting in 4% agarose and submerged in culture media. Hearts were imaged on an Olympus IXplore SpinSR system with a Yokogawa CSU-W1 SoRa spinning disk unit and 30x/1.05NA silicone immersion objective. Imaging was performed in an environmental chamber at 28 °C without CO_2_. Z stacks were acquired every 2.5 minutes for 30-100 minutes. Images and videos were processed using Fiji.

### Photoconversion of Kaede larvae

To photoconvert Kaede, *TgBAC(csf1ra:Gal4i); Tg(UAS:kaede)* embryos at 48 hpf were anaesthetised and embedded ventrally in 1% low melting point agarose. A region of interest (ROI) was set around the heart and a z-stack through the entire cardiac region was set. Photoconversion was performed by illuminating the selected ROI with a 405 nm laser at 100% power for the duration of the z-stack. Embryos were recovered and their hearts imaged at specified time points following heart dissection.

### Fluorescence associated cell sorting (FACS)

Dissected hearts were gently torn open and washed in ice-cold perfusion buffer (10 mM HEPES, 30 mM Taurine, 5.5 mM Glucose in PBS). Tissue was digested in perfusion buffer plus 5 mg/ml Collagenase II (Worthington Biochemical Corp; LS004176) for up to 2 hours at 32 °C, agitated at 800 rpm, with regular pipetting to aid digestion. To stop digestion, the cell suspension was placed on ice and mixed with one volume of perfusion buffer plus 10% (vol/vol) FBS (Fisher Scientific; 11591821). Cells were pelleted (1500 rpm (7500 x g), 5 minutes, 4°C), resuspended in PBS plus 2% FBS and filtered through a 40 μm Falcon® cell strainer (Fisher Scientific; 22363547).

All cells were kept on ice until analysis. Immediately prior to sorting, DRAQ7TM (Abcam; ab109202) was added to the cell suspension to stain dead cells. All flow cytometry analysis and FACS was carried out at 4°C using a BD Influx Fluorescence Associated Cell Sorter or a BD FACS ARIA™ II SORP Flow Cytometer Cell Sorter. For sorting and analysis, events were gated to select single, live cells. Data were analysed using FlowJo_v10.6.2 software (BD Life Sciences).

### Cytology

FACS cells were resuspended in 200 μl PBS and loaded into Shandon™ EZ Single Cytofunnels (Thermo Scientific; 11972345) and spun (1000 rpm, 5 minutes) onto coated Shandon Cytoslides (Thermo Scientific; 12026689) using a Cytospin Cytocentrifuge (Thermo Scientific). Slides were dried and fixed in 100% methanol for 5 minutes, stained in May-Grünwald’s stain for 5 minutes, then transferred to Giemsa stain for 10 minutes. Slides were rinsed in 3.3 mM Sorenson buffer (pH 6.8) for 3 minutes to differentiate then left to dry. Cytospins were imaged on an Olympus BX53 Upright Microscope with a 40X dry objective.

### Image Analysis and statistics

Images and videos were processed using Fiji^57^. For manual cell counting, images were blinded and counted using the Cell Counter plugin in Fiji (https://imagej.nih.gov/ij/plugins/cell-counter.html).

For the automated cell shape analysis, a freely available Modular Image Analysis (MIA; version 0.21.8) workflow automation plugin for Fiji was used^58^. Briefly, cells were detected in both channels using the same method: 2D median filter applied to reduce image noise whilst retaining sharp object edges; image binarised using global Otsu threshold^59^, and an applied threshold added. The binarised image was subject to distance-based watershed transformation and 2D holes (enclosed background pixel regions) filled. Objects were detected as contiguous regions of foreground-labelled pixels^60^; detected cells subject to minimum volume threshold. Any cells smaller than this threshold were discarded from further analysis. Cells in contact with the 2D image edge (i.e. not considering Z-axis) were also removed from further analysis as their areas could not be accurately determined. Cells from channels 1 and 2 were related based on their mutual overlap. Any cells with at least 25% overlap with another cell from the opposite channel were classed as being related. The number of related (overlapping) cells was calculated. 2D ellipses were fit to each cell and used to calculate eccentricities^61^.

All data was recorded and analysed using Excel and Graphpad Prism (v10). All data was checked for outliers via a Grubb’s test (alpha = 0.05) and any outliers removed. All data was then checked for normal distribution via a Shapiro-Wilk test (alpha = 0.05) and parametric or non-parametric tests chosen accordingly. Details of statistical tests used are provided in figure legends. Wherever possible the researcher was blinded to the status of control/test subjects via the use of individual identifier codes for each animal.

### 3D image rendering

3D projections of adult hearts were generated on IMARIS software using maximum intensity projections of specified transgenic lines.

### 3D tracking of macrophages

For the tracking of cells in *xyzt* from the *ex vivo* imaging platform, a plugin within MIA (version 0.21.8) was used. The plugin segments cells detected in two channels and tracks their migration in 3D.

## Results

### Larval zebrafish hearts are populated by embryonic macrophages

Adult zebrafish are known to have populations of cTMs in the steady state and cardiac MNP populations expand following cardiac injury^24,29,62^. A recent study has highlighted the heterogeneity of resident cTMs and their involvement in promoting regeneration^42^ echoing what has been described in neonatal mice^63^. However, we are yet to fully understand the spatial distribution of these cells in the heart, the roles played by distinct MNP lineages and to confirm strategies to identify these subpopulations. We aimed, therefore, to understand the dynamics of distinct cardiac MNPs in the zebrafish to determine whether differences in ontogeny and regulation of their inflammatory profile contribute to regeneration. Small numbers of *mpeg1.1*+ cells have previously been noted in the uninjured larval heart and in the pericardium^64^, but the precise timing of cTM recruitment has not been investigated. Initially we evaluated when putative resident macrophages colonise the heart during zebrafish development using two widely used macrophage transgenic marker lines: *mpeg1.1* and *csf1ra*. We imaged hearts from Tg(*myl7:GFP*); Tg(*mpeg1.1:mCherry*) double transgenic fish at 2-24 dpf and assessed the number of cells and their distribution in the cardiac chambers (Figure 1A-C). Very small numbers of *mpeg1.1*+ cells were observed in the heart at 2-7 dpf (2 dpf mean = 0, 3 dpf mean = 0.4 ± 0.7 SD, 5 dpf = 0.4 ± 0.5 SD, 7 dpf = 0.9 ± 0.9) with slight but non-significant increases observed at 10 (1.9 ± 1.6 SD) and 14 dpf (5.4 ± 3.7 SD). At 17 dpf significant numbers of *mpeg1.1*+ cells were observed in the heart (11.7 ± 5.3 SD) and numbers increased from there on (Figure 1A and B). Interestingly, when we repeated this experiment using TgBAC(*csf1ra:GFP*); Tg(*myl7:mCherry*) double transgenic fish we observed significantly more *csf1ra*+ cells than *mpeg1.1*+ cells at 3, 5, 7 and 10 dpf (Figure 1A-C). The number of *csf1ra*+ cells remained constant over the entire timecourse whereas the number of *mpeg1.1*+ cells increased markedly after 10 dpf (Figure 1B).

**Figure 1.**
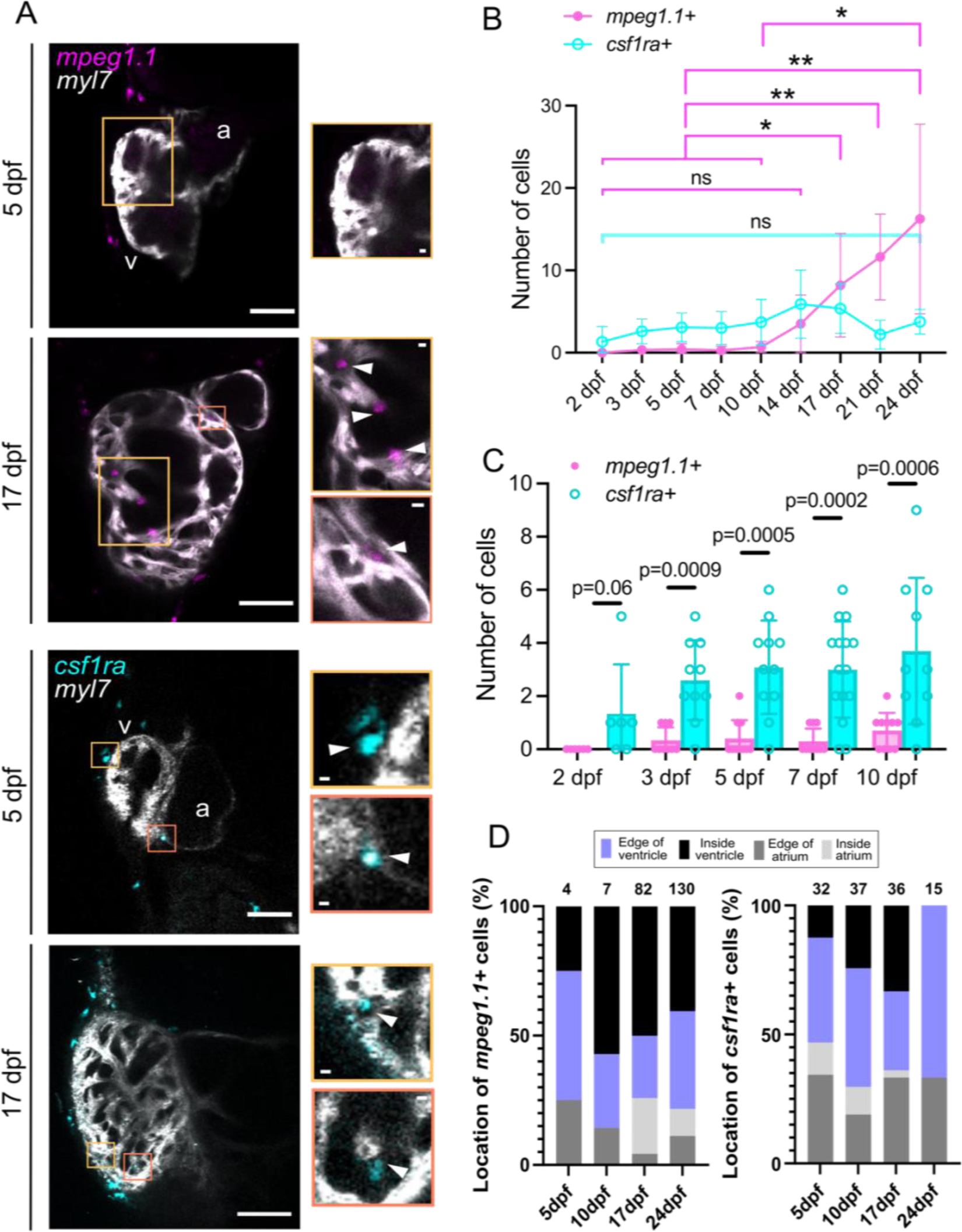
The zebrafish heart is seeded with macrophages during embryonic stages. **A**) Confocal maximum projections of Tg*(myl7:GFP);* Tg*(mpeg1.1:mCherry)* (top) and Tg*(myl7:mCherry);* TgBAC*(csf1ra:GFP)* (bottom) double transgenic hearts at 5 and 17 dpf. In both cases the myocardium (*myl7*) is shown in grey. *mpeg1.1*+ cells are in magenta and *csf1ra*+ cells in cyan. Colour-coded boxed regions indicate the position of insets. White arrowheads in insets show location of *mpeg1.1* and csf1ra-positive macrophages. v = ventricle, a = atrium. Scale bars = 25 μm and 5 μm for insets. **B,C**) Graphs indicating the number of *csf1ra±* and *mpeg1.1±* cells in larval hearts from (B) 2-24 dpf and (C) 2-10 dpf (same data). N = 6-10 larvae per timepoint. Statistical analysis in B: Kruskal-Wallis with Dunn’s multiple comparisons test for each cell population timecourse separately. Statistical analysis in C: Mann-Whitney test between the two cell populations at each timepoint. Ns = not significant. **D**) Bar charts showing the proportion of *mpeg1.1+* (left) and *csf1ra+* (right) cells in the ventricle and atrium at the timepoints indicated. The distribution of cells within each chamber is also indicated. Numbers above bars indicate total macrophage number counted. N = 5-12 larvae per timepoint.

When assessing the tissue distribution of these cells in the larval and juvenile heart, we observed different localisation for the two populations of cells. Throughout the timecourse *mpeg1.1*+ cells were more prevalent in the ventricle than the atrium (78.3% ± 4% SD in the ventricle at 24 dpf; Figure 1D). By contrast, *csf1ra+* cells were evenly distributed between the ventricle and atrium at 5 dpf, and although the majority were present in the ventricle at later stages (66.6% ± 4.5% SD at 24 dpf), a higher proportion were found in the atrium compared to *mpeg1.1+* cells (Figure 1D). We also observed that *csf1ra*+ cells were progressively found only on the surface of the atrium and ventricle whereas *mpeg1.1*+ cells were initially located on the surface but were later found distributed throughout the tissue (Figure 1D).

Collectively, our data suggests that the zebrafish heart is first seeded with *csf1ra*+ embryonic cTMs during early development with additional *mpeg1.1*+ cells observed at later larval stages. Additionally, although *csf1ra*+ and *mpeg1.1*+ have both been suggested to label primitive and definitive haematopoiesis-derived embryonic macrophages^13,36,39,65^, our findings suggest they may indicate different subpopulations during cardiac development.

### Embryonic cTMs are primitive haematopoiesis-derived and retained into juvenile stages

As the observed expansion of *mpeg1.1*+ macrophages in the heart corresponds to the reported timing for the onset of HSC-derived haematopoiesis^65,66^, we next aimed to determine the origin of zebrafish cardiac macrophages. First, we investigated if the embryonic-derived *csf1ra+* cTMs are retained in the heart beyond 10-14 dpf, when HSCs begin to contribute significantly to myeloid cell lineages^66,67^. Using the TgBAC(*csf1ra:Gal4i);* Tg(*UAS:kaede*) line we photoconverted all the cTMs present at 2 dpf and examined the hearts of these fish at 21 dpf (Figure 2A,B). Photoconverted (yellow) cTMs were still observed in the heart at 21 dpf, although larger numbers of non-photoconverted (cyan) *csf1ra*+ cells were also present (Figure 2A,B). This suggests that the early seeded embryonic *csf1ra*+ cTMs are retained during late larval stages of development following the switch to HSC-derived haematopoiesis, although additional cells may contribute to cTMs after 2 dpf.

**Figure 2.**
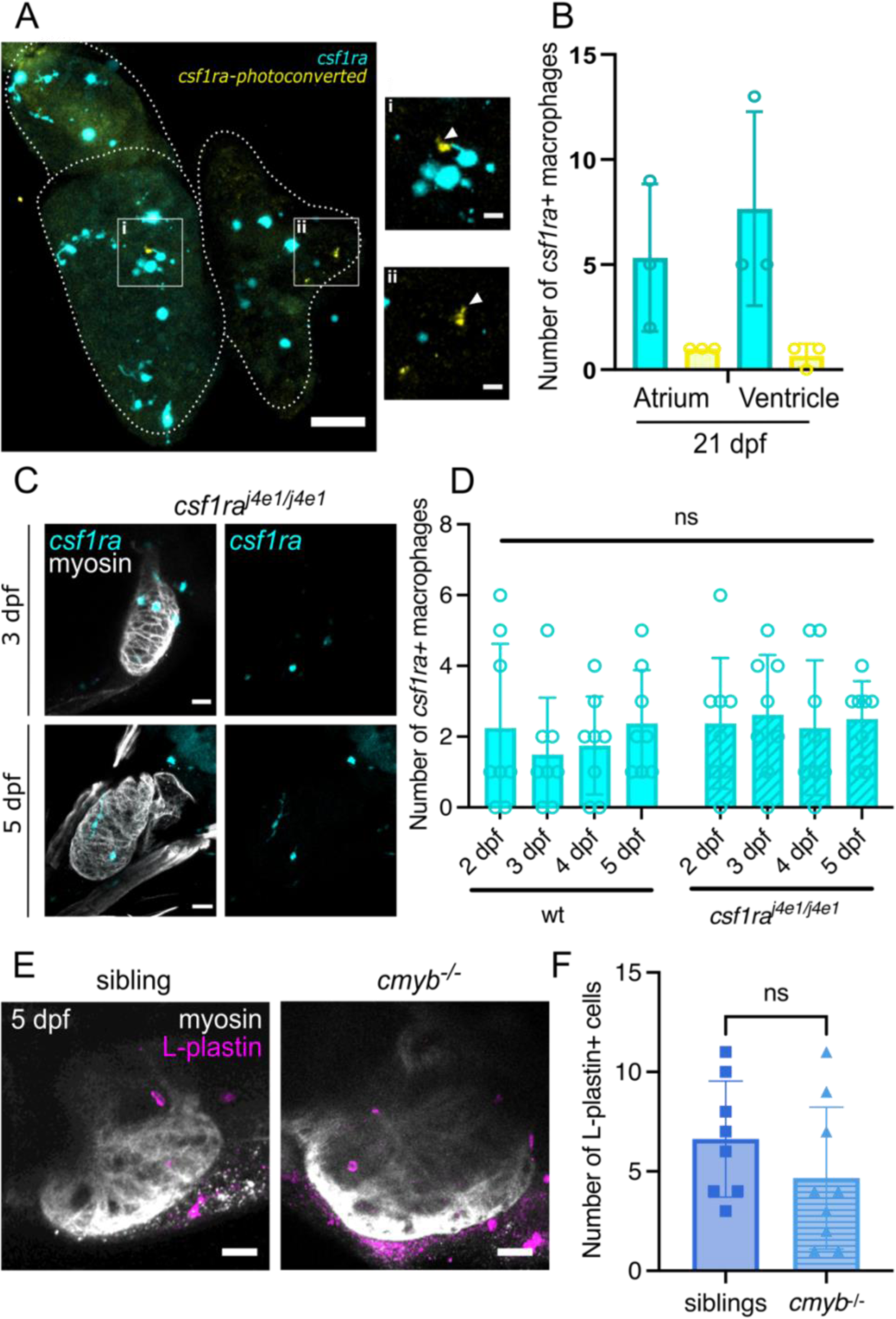
Embryonic cTMs are retained at juvenile stages and are primitive haematopoiesis derived. **A**) Confocal maximum projection image of a Tg*(csf1ra:Gal4i));* Tg*(UAS:kaede)* larval heart at 21 dpf after photoconversion at 2 dpf. Scale bars = 50 μm Zoomed insets show photoconverted (yellow) *csf1ra*-positive cells in the i) ventricle and ii) atrium. Scale bars = 10 μm. **B**) Quantification of *csf1ra* unphotoconverted (cyan) and photoconverted (yellow) macrophages at 21 dpf, n = 3. **C**) Maximum projection images of *csf1ra^j4e1/j4e1^*; *TgBAC(csf1ra:GFP)* larval hearts at 3 and 5 dpf. An anti-myosin antibody was used to label the myocardium. Scale bars = 25 µm. **D**) Graph showing quantification of *csf1ra* macrophage number in hearts of wildtype and *csf1ra^j4e1/j4e1^*; *TgBAC(csf1ra:GFP)* hearts at the time-points indicated. N = 8 per genotype. Statistical analysis = Welch’s t-tests between genotype per timepoint. (**E,F**) Maximum projection images (E) and quantification (F) of the number of L-plastin+ cells in the heart of *cmyb^−/−^* larvae and their siblings (het/wt) at 5 dpf. N = 9/8. Statistical analysis = Welch’s t-test.

To further determine the ontogeny of these embryonic and larval seeding cTMs, we assessed the number of macrophages present in the hearts of mutant lines that affect different haematopoietic lineages. Emergence of primitive macrophages has been shown to be normal in *csf1ra* mutant zebrafish^13,65,68^. Mutants exhibit defective microglia formation, but normal seeding of peripheral tissues with all macrophages suggested to be primitive-derived up to 5 dpf^13,65,68^. We, therefore, investigated the embryonic seeding of cardiac macrophages in *csf1ra^j4e1/j4e1^*; Tg*(mpeg1.1:mCherry);* TgBAC(*csf1ra:GFP)* fish during development (Figure 2C,D). No differences in the numbers of cTMs were observed in wildtype and *csf1ra^j4e1/j4e1^*fish at 2-5 dpf, suggesting that these early *csf1ra+* cTMs are derived from primitive haematopoiesis. Similarly, no significant difference in immune cell number was observed in the hearts of *cmyb^−/−^* mutants, which have normal primitive haematopoiesis but lack HSCs^49,67^, at 5 dpf (Figure 2E,F) further suggesting that the earliest cTMs in the heart are derived from primitive haematopoiesis.

### Adult zebrafish hearts contain subpopulations of Csf1ra-expressing cells

We have previously analysed co-expression of the two commonly used macrophage markers, *csf1ra* and *mpeg1.1*, in the adult zebrafish heart, where we identified a population of *mpeg1.1*+ lymphocytes^69^. To further characterise the precise distribution of different MNP (and lymphoid) populations in the unwounded zebrafish heart, confocal imaging of hearts from transgenic adult zebrafish was performed (Figure 3). Imaris 3D renders of *mpeg1.1*+ MNPs in Tg(*mpeg1.1:GFP*); Tg(*myl7:HRAS-mCherry*) (Figure 3A, Supplementary video 1) and Tg(*mpeg1.1:GFP*); Tg(*kdrl:mCherry-CAAX*) fish (Figure 3B, Supplementary video 2) highlight the different morphologies and distribution of these cells on the surface of the heart. Rounded cells could be observed embedded within, and in direct contact with, several cardiomyocytes (asterisks in Figure 3A, Supplementary video 1) and more elongated cells were observed close to cardiac vessels (Figure 3B, Supplementary video 2). Next, we analysed imaging data from TgBAC(*csf1ra:GFP*); Tg(*mpeg1.1:mCherry*) adult hearts, and determined cell size and shape in 3D using a custom Fiji plugin^58^. The majority of labelled cells in the ventricle expressed both transgenes (*mpeg1.1+ csf1ra+* cells), were distributed across the surface of the heart and possessed a stellate morphology, typical of cardiac tissue macrophages (Figure 3A-C)^15,24^. Cells expressing only *csf1ra:GFP,* with a similar elongated morphology (*csf1ra*+ cells), were also observed as were the rounded cells expressing only *mpeg1.1:mCherry* (*mpeg1.1*+ cells) we have described previously^69^ (Figure 3A-C). Analysis of fluorescence and shape of cells on the surface of the ventricle showed that 49.5% of labelled cells were *mpeg1.1+ csf1ra+*, 18.2% *csf1ra*+ and 32.2% were *mpeg1.1*+. No differences in size or circularity between *mpeg1.1+ csf1ra+* and *csf1ra*+ cells were observed but *mpeg1.1*+ cells were significantly smaller and rounder than *csf1ra*-expressing cells, as previously described for 2D analysis (Figure 3E)^69^. Analysis of the number of cellular projections revealed that *mpeg1.1+ csf1ra+* cells were significantly more protrusive than both *mpeg1.1*+ and *csf1ra*+ cells, suggesting functional differences between the two *csf1ra*-expressing populations (Figure 3D). Similar distribution of cells within the atrium was observed, although there was a higher proportion of *mpeg1.1*+ lymphocytes (34.4% of labelled cells were *mpeg1.1+ csf1ra+*, 11.0% *csf1ra*+ and 54.5% *mpeg1.1*+) (Figure 3C). These data indicate that significant numbers of three populations of immune cells can be distinguished, in both the ventricle and the atrium, via these two transgenic lines.

**Figure 3.**
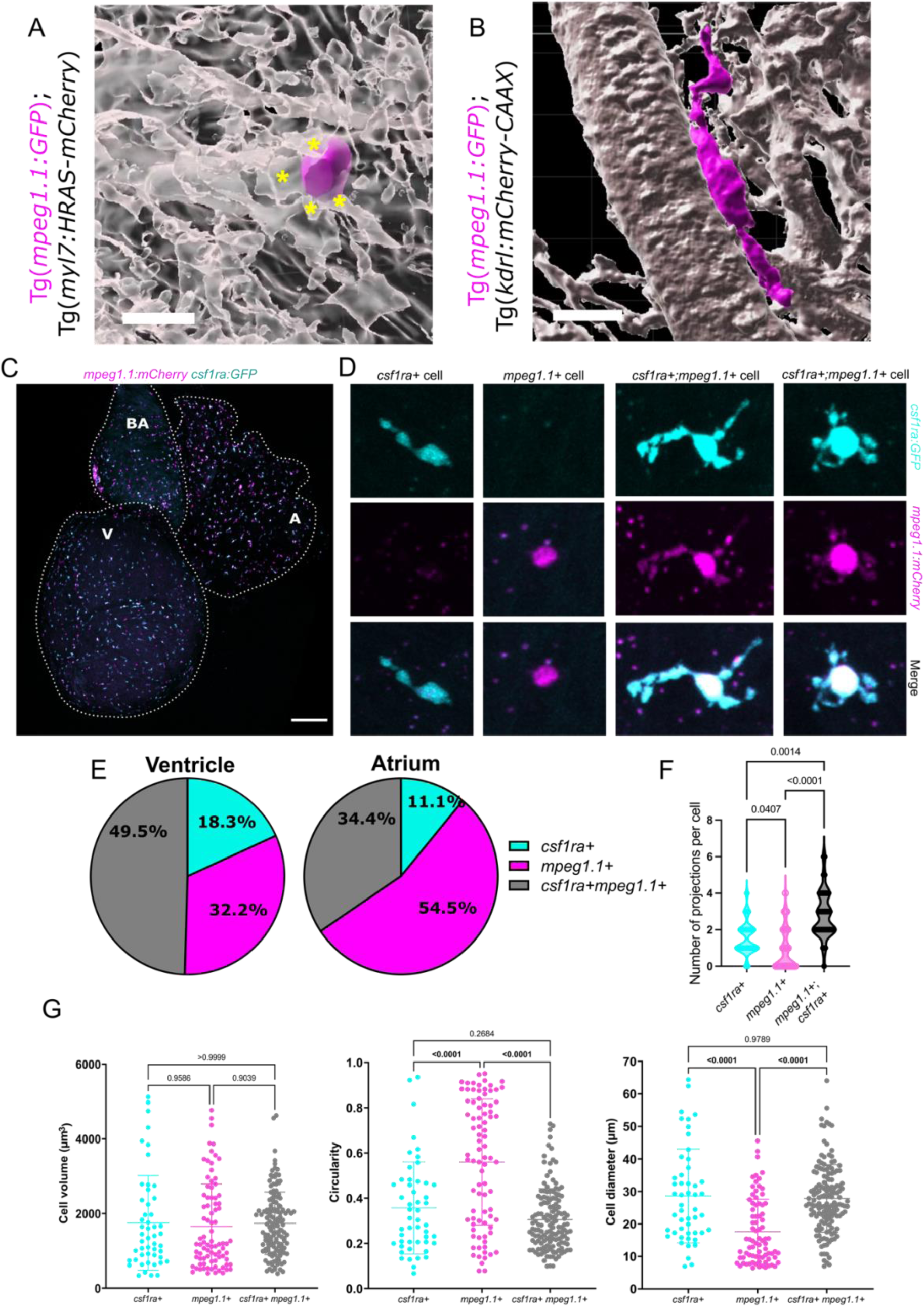
Subpopulations of MNPs can be identified in the adult zebrafish heart via Tg*(mpeg1.1:mCherry);* TgBAC*(csf1ra:GFP)* transgenic lines. **A,B**) 3D still views of Imaris renders (taken from supplementary videos 1 & 2, respectively) of individual *mpeg1.1*+ (magenta) cells in the ventricle associated with *myl7*+ cardiomyocytes (A) or *kdrl*+ endothelial cells (B; both shown in grey). Yellow asterisks in A demark individual CMs in direct contact with the *mpeg1.1*+ macrophage. Scale bar = 10 µm. **C**) Wholemount imaging of a Tg*(mpeg1.1:mCherry)*; TgBAC*(csf1ra:GFP)* double transgenic adult heart. The dotted line demarcates the ventricle (V), atrium (A) and bulbus arteriosus (BA). Scale bar = 200 μm. **D**) High magnification views of individual cells from the ventricle showing differing cell morphology between subtypes. **E**) Graphical representation of the relative proportions of labelled cells found in the ventricle and atrium. **F**) Number of cellular projections per cell type. Statistical analysis performed using a Kruskal-Wallis test. Projections counted manually from 28-46 cells per cell type across three ventricles. **G**) Automated cell shape analysis of *mpeg1.1± csf1ra±* cells from a 388 µm^2^ Z-stack imaged within the ventricle. Cells were classified by their expression of *mpeg1.1:mCherry* and/or *csf1ra:GFP* and individual cell volume, circularity, and diameter was measured. Statistical analysis was performed by Brown-Forsythe and Welch’s ANOVA test following removal of outliers by the ROUT outlier test (Q = 1%). 50-147 cells measured across three hearts.

### Macrophages are differently distributed within the ventricle and specific subpopulations are defective in Csf1ra mutants

As both *csf1ra*+ and *mpeg1.1+ csf1ra+* populations of macrophages could be observed on the surface of adult zebrafish hearts we aimed to determine their distribution throughout the cardiac tissue. It has been reported that embryonically derived CCR2-cTMs have a different distribution to monocyte-derived populations in rodents^19^, but this has not been assessed in zebrafish. Therefore, to further assess the distribution of different MNP (and lymphoid) populations within the entire heart, we employed tissue clearing on double transgenic TgBAC(*csf1ra:GFP*); Tg(*mpeg1.1:mCherry*) fish (Figure 4). Ce3D tissue clearing^54^ revealed cells at all depths through the ventricle (Figure 4A-C; Supplementary video 3). Quantification of *mpeg1.1+ csf1ra+*, *csf1ra*+ and *mpeg1.1*+ cells at different depths revealed differences in their distribution (Figure 4C). Large numbers of *mpeg1.1+ csf1ra+* cells were observed at all depths, with a trend towards more on the surface of the heart (Figure 4C). Conversely, *mpeg1.1*+ cells were evenly distributed throughout the ventricular wall (Figure 4C). Strikingly, *csf1ra*+ cells were only observed directly on the surface of the heart and were absent from the deeper tissue (Figure 4B,C), the same pattern observed for embryonically derived cTMs in the rodent heart^19^.

**Figure 4.**
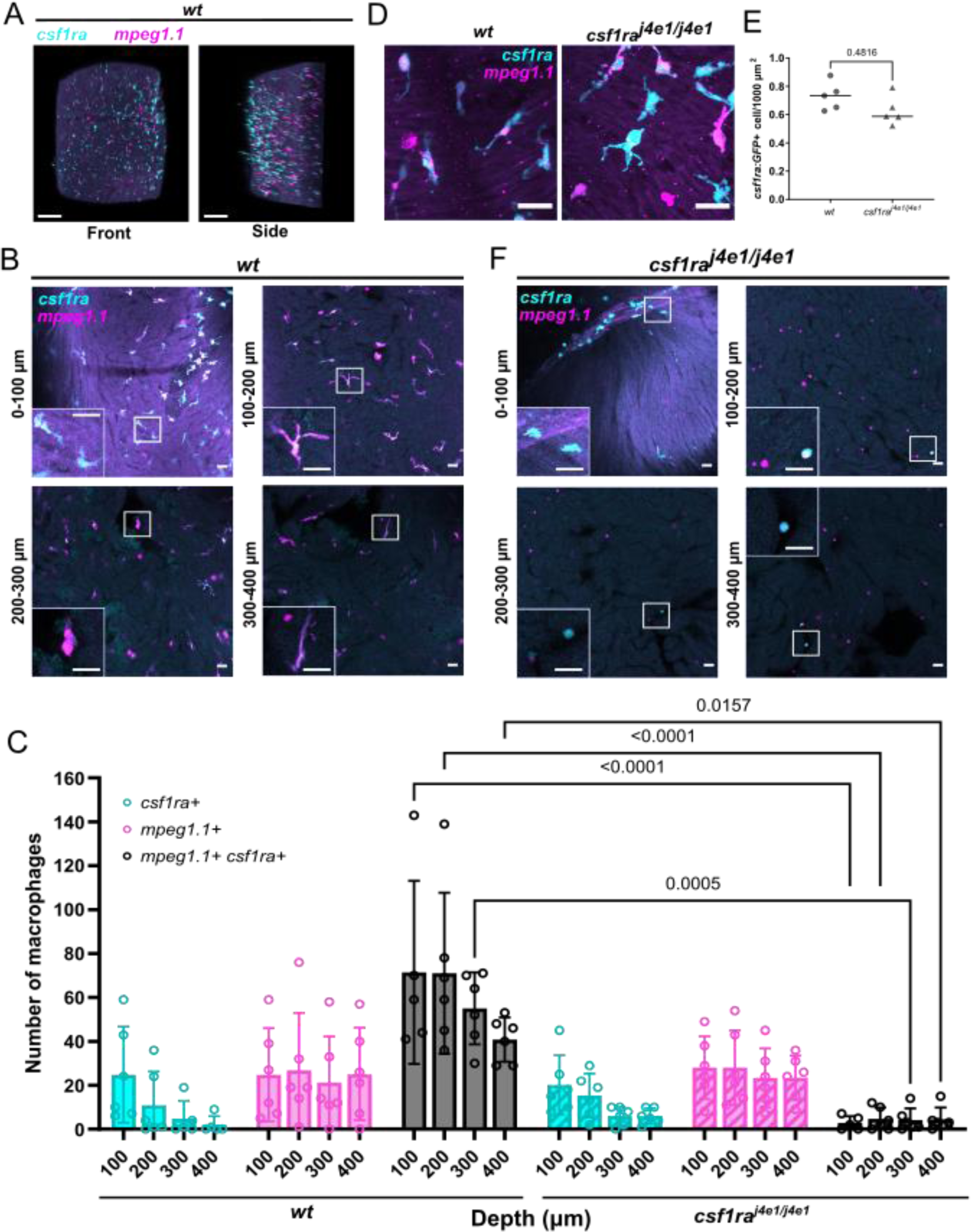
Analysis of immune cell distribution in wildtype and *csf1ra^j4e1/j4e1^*; Tg(*mpeg1.1:mCherry*); TgBAC(*csf1ra:GFP*) adult hearts. **A**) 3D reconstruction of a cleared adult wildtype heart fluorescently stained with anti-GFP (*csf1ra*) and anti-mCherry *(mpeg1.1*) antibodies showing front and side views. Scale bars = 100 µm. **B**) Single, representative z-slices from cleared Tg(*mpeg1.1:mCherry*); TgBAC(*csf1ra:GFP*) ventricles at specified depths (0-400 µm). Insets show enlarged regions of boxed cells. Scale bars (main and inset) = 20 µm. **C**) Graph indicating mean frequency ± SD of *csf1ra:GFP± mpeg1.1:mCherry*± populations per 100 µm sub-stack depth through the ventricle of wildtype and *csf1ra^j4e1/j4e1^*; Tg(*mpeg1.1:mCherry*); TgBAC(*csf1ra:GFP*) fish. The Z-stack was divided into 200 x 200 x 100 µm (xyz) sub-stacks and *csf1ra:GFP± mpeg1.1:mCherry*± cells were manually counted using the Cell Counter plugin on Fiji following blinding of datasets. All cells were counted per field of view, n= 6 per genotype. Statistical analysis was performed on mean frequency per fish by two-way ANOVA with Šidák’s multiple comparisons test. **D**) Representative images of fluorescent cells on the ventricular surface of wildtype and and *csf1ra^j4e1/j4e1^*; Tg(*mpeg1.1:mCherry*); TgBAC(*csf1ra:GFP*) hearts. **E**) Automated quantification of *csf1ra:GFP+* cell number per 1000 µm^2^ of the ventricle surface. N = 5 per genotype. Statistical analysis was performed by Kruskal-Wallis test. **F**) As in (B), single, representative z-slices from *csf1ra^j4e1/j4e1^*; Tg(*mpeg1.1:mCherry*); TgBAC(*csf1ra:GFP*) mutant ventricles at specified depths. Insets show enlarged regions of boxed cells. Scale bar = 20 µm.

We have previously shown that *csf1ra^j4e1/j4e1^* fish exhibit a significant reduction in *mpeg1.1+ csf1ra+* cells via FACS, although *csf1ra*+ and *mpeg1.1*+ numbers appeared normal^69^. We, therefore, aimed to determine the distribution of labelled cells in *csf1ra^j4e1/j4e1^*; Tg*(mpeg1.1:mCherry);* TgBAC(*csf1ra:GFP)* ventricles (Figure 4C-E). Surprisingly, whole mount imaging revealed *csf1ra*+ and *mpeg1.1*+ *csf1ra*+ cells with a typical stellate morphology on the surface of the ventricle of all fish examined (Figure 4D). Indeed, quantification of the number of surface *csf1ra*:GFP-expressing MNPs revealed no differences between wildtype and *csf1ra^j4e1/j4e1^* fish (Figure 4E). Loss of *csf1ra* has been suggested to affect cell dispersal in the adult zebrafish brain^71^, so we next assessed the distribution of cTMs within the deep myocardium via Ce3D clearing. As observed for confocal surface imaging, *csf1ra*+ cells were present mainly on the surface of *csf1ra^j4e1/j4e1^*ventricles (Figure 4C,F), however, strikingly, *mpeg1.1+ csf1ra+* MNPs were markedly diminished at all depths (Figure 4C,F; Supplementary video 4).

### Zebrafish have orthologs to mammalian MNP markers

In an effort to further assess conservation of mammalian MNP markers in zebrafish, we summarised the predicted presence or absence of zebrafish orthologs to mouse genes which are routinely used to define populations of MNPs. Through literature searches, genes of interest were compiled (Supplementary Table S1A) and investigated using the Alliance of Genome Resources database (The Alliance; Version 5.0.0; https://www.alliancegenome.org/). The Alliance compiles genetic information from seven model organism databases and the Geno Ontology Consortium and provides orthology predictions using the DRSC Integrative Ortholog Prediction Tool (DIOPT). The DIOPT integrates multiple orthology databases and summarises the gene orthology predicted by these databases for each species (Supplementary Table 1A). This confirmed that Csf1r signalling is highly conserved in zebrafish, as *csf1ra* was detected to be the main ortholog to murine *Csf1r* in 10/11 databases (Supplementary Table 1A). The paralog of *csf1ra*, *csf1rb,* was only detected in 2/11 databases. Interestingly, orthologs of monocyte markers *Ly6c1* and *Sell/Cd62l* were not found in the zebrafish genome. Apart from these genes, orthologues to all other MNP markers investigated were found in the zebrafish genome, although some were only predicted by a small number of databases (Supplementary Table 1A). For example, zebrafish *ccr2* was detected to be the *Ccr2* ortholog in 5/11 databases, but interestingly, was also detected to be the only potential ortholog of *Cx3cr1*, although this was only predicted by one database. This conservation of *ccr2* is particularly interesting considering the use of this cell surface receptor to identify monocytes/MDMs that have been repeatedly shown to elicit negative effects on tissue repair ^20,72,73^ and suggests that analogous populations may be present in zebrafish.

We also performed the reverse orthology analysis on commonly used zebrafish genes and showed that human and mouse had commonly identified orthologs to *marco*, *mfap4*, *apoeb*, *acod1* (*irg1*), *mhc2dab* and *mertka* (Supplementary Table 1B). Interestingly, *MPEG1* and *Mpeg1* were only predicted to be an ortholog to *mpeg1.1* in 2/11 databases but were predicted to be *mpeg1.2* in 10/11 databases. Accordingly, gene expression analysis showed *mpeg1.2* expression in all cardiac *csf1ra*-expressing MNPs (see Figure 5), but lacking from the *mpeg1.1*+ lymphoid population^69^. It is likely, therefore, that *mpeg1.2* may be a superior marker for MNPs than *mpeg1.1*. We also assessed other commonly used lineage specific genes to further clarify their expression to hematopoietic lineages, such as markers for T cells, NK cells, and neutrophils ^74–76^. Collectively, this analysis confirmed the high degree of evolutionary conservation of genes utilised by mammals and zebrafish for MNP homeostasis albeit with some notable exceptions. These sub-population stratification markers warrant further investigation in zebrafish.

**Figure 5.**
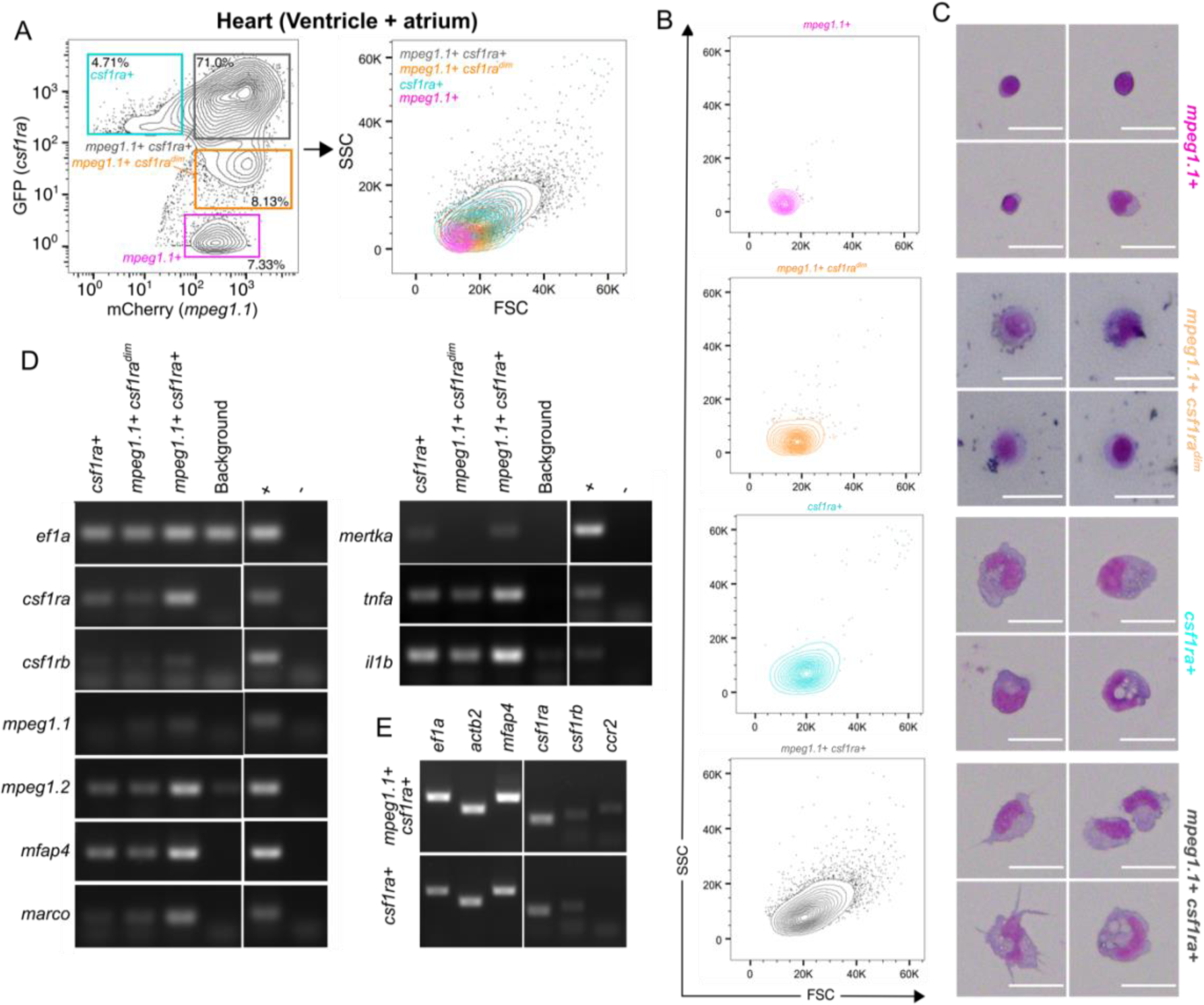
*csf1ra*+ MNPs exhibit differing morphologies and marker expression. **A,B**) Flow cytometry plot of cell populations gated by GFP and mCherry expression from Tg*(mpeg1.1:mCherry)*; TgBAC*(csf1ra:GFP)* hearts (left plot) and corresponding FSC and SSC of these populations (all gates merged in right plot, individual populations shown in B). **C)** May-Grünwald Giemsa staining of sorted *csf1ra±* and *mpeg1.1±* sub-populations. Cells were isolated from 6-8 pooled Tg*(mpeg1.1:mCherry)*; TgBAC*(csf1ra:GFP)* hearts. Scale bar = 20 µm. **D**) RT-PCR analysis of sorted populations using MNP related markers. cDNA generated from whole kidney marrow was used as a positive control; water was used for the negative control. Cells were sorted from pools of 15 hearts for each group. Cell numbers: *csf1ra*+ = 3300; *mpeg1.1*+*csf1ra*^dim^ = 4900; *mpeg1.1*+ *csf1ra+* = 30,000; background cells = 250,000. **E**) RT-PCR analysis of MNP markers from sorted cells isolated from 1 dpi Tg(*mpeg1.1:mCherry*); TgBAC(*csf1ra:GFP*) ventricles. Cell counts: mpeg1.1+; csf1ra+ = 7900; csf1ra+ = 712.

### *csf1ra* and *mpeg1.1* double transgenic zebrafish reveal four immune cell populations in cardiac tissue

We next used our generated orthologous marker resource to interrogate the identity of the different *mpeg1.1*+ and *csf1ra*+ cell populations in the adult zebrafish heart. In order to determine markers of each of the identified populations, we sorted large numbers of whole hearts (including ventricle and atrium) from adult TgBAC(*csf1ra:GFP*); Tg(*mpeg1.1:mCherry*) fish (Figure 5). In doing so, we identified an additional double positive population that were dim for GFP (*mpeg1.1+ csf1ra^dim^*; Figure 5A). Analysis of forward and side scatter profiles of each population suggests the *mpeg1.1+ csf1ra^dim^* cells are smaller and less granular than *csf1ra*+ and *mpeg1.1+ csf1ra*+ cells although all *csf1ra*+ cells remain larger than the *mpeg1.1+* lymphoid cells, as previously described^69^ (Figure 5B). Cytology further confirms these findings (Figure 5C). We next assessed the expression of the panel of conserved myeloid markers across the three *csf1ra*+ populations via RT-PCR (Figure 5D). Interestingly, all markers were found in each population, in varying amounts, with the exception of *mertka*, which was absent from the *mpeg1.1+ csf1ra^dim^* population (Figure 5D). The *mpeg1.1+ csf1ra^dim^*population also expressed lower levels of the macrophage differentiation markers *mmd*, *mfap4* and *marco* and of pro-inflammatory markers *tnfa* and *il1b* (Figure 5D). *csf1ra*+ cells also expressed lower levels of these markers than *mpeg1.1+ csf1ra*+ cells (Figure 5D).

As *CCR2* expression has been used to distinguish monocyte-derived macrophages from tissue resident populations in the mouse model, we also analysed this marker in the different populations of cells. Expression of *ccr2* was difficult to detect in cells sorted from uninjured hearts but when sorting cells from 1 day post cryoinjury (dpi) hearts, *ccr2* was detectable in *mpeg1.1+ csf1ra*+ cells but was absent from *csf1ra*+ cells (Figure 5E). This further suggests that the *csf1ra*+ cells found on the surface of the adult heart could be an embryonic-derived tissue resident population of macrophages.

### Cardiac MNP populations exhibit different behaviours

To further understand the behaviour of the different MNPs within the intact heart, we sought to visualise their movement in real time. Adult zebrafish hearts are difficult to image *in situ* due to their depth within the body and the opaque nature of the surrounding tissue. Methods for *ex vivo* culture of whole zebrafish hearts have recently been described and used to study regenerative processes including revascularisation, cardiomyocyte proliferation and epicardial activation^55,77,78^. However, *ex vivo* culture has not yet been used to visualise macrophage movement within intact cardiac tissue. We modified existing protocols^55^ and imaged uninjured hearts from Tg(*mpeg1.1*;*mCherry*); TgBAC(*csf1ra*:*GFP*) zebrafish (Figure 6A). As in fixed tissue, *csf1ra*+ and *mpeg1.1*+ *csf1ra*+ myeloid cells and *mpeg1.1*+ lymphoid cells were observed on the surface of the heart and all showed movement (Figure B-D; Supplemental video 5). The behaviour of *mpeg1.1*+ *csf1ra*+ and *csf1ra*+ myeloid cells was compared and classified into groups (Figure 6D). Both cell populations were observed to only rarely be completely static or to appear into the imaging plane from deeper tissues. A few cells disappeared out of the imaging plane, suggesting they were migrating deeper into the tissue (Figure 6D). The majority of cells were observed to either be stationary but forming constant protrusions or to be actively traversing within the imaging plane e.g. on the surface of the heart (Figure 6B,C; Supplemental video 5). Interestingly, the two macrophage populations showed differing behaviours with *csf1ra*+ cells being more likely to traverse than *mpeg1.1*+ *csf1ra*+ cells but less likely to be stationary and protruding (Figure 6B-D; Supplemental video 5). This data indicates that the two *csf1ra*+ populations exhibit different behaviours and these can be revealed by *ex vivo* imaging of the heart.

**Figure 6.**
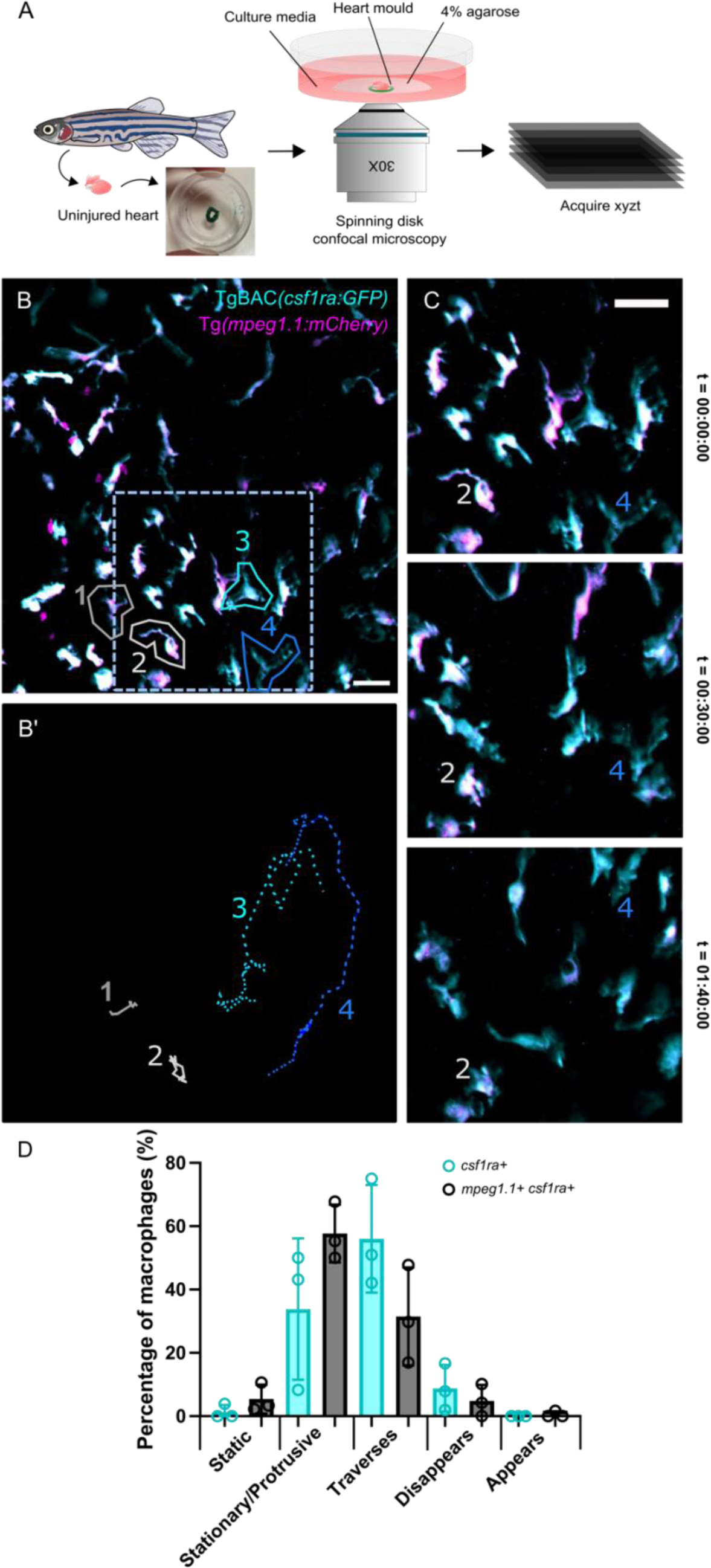
*Ex vivo* live imaging of cardiac *csf1ra±* and *mpeg1.1±* cells reveals different behaviours. **A**) Schematic of the *ex vivo* imaging platform. Single hearts were placed in a custom mould (green) and mounted in 4% low-gelling agarose to limit transverse movement of the heart during imaging. **B**) Maximum projection of *csf1ra±* and *mpeg1.1±* positive cells on the surface of the ventricle. Two *mpeg1.1+ csf1ra+* (1/2; grey) cells and two *csf1ra*+ cells (3/4; cyan/blue) are highlighted. Scale bar = 20 µm. **B’**) Colour coded tracks of cells 1-4 from B. Cells were tracked from 0 – 100 minutes using the Manual Cell Tracking Plugin on Fiji. **C**) Maximum projection images of the boxed region in B at the timepoints indicated. Cells 2 (*mpeg1.1+, csf1ra+*) and 4 (*csf1ra*+) are indicated in each frame. Scale bar = 20 µm. **D**) Quantification of *mpeg1.1+ csf1ra+ and csf1ra+* MNP behaviour. n = 3 (total cell number analysed: *csf1ra+* = 12-51; *mpeg1.1+ csf1ra+* = 47-90).

### cTMs proliferate after injury

Finally, we aimed to determine how the identified macrophage populations respond to injury. We and others have shown that there is a rapid and robust inflammatory response to cardiac damage^24,29,62^ and that this comprises different macrophage inflammatory states^24^. However, how different macrophage ontogenies respond to injury is not well defined. We have previously shown that *csf1ra^j4e1/j4e1^* fish exhibit reduced pro-inflammatory macrophage activation and reduced collagen deposition following cardiac cryoinjury^24^. We, therefore, analysed hearts from Tg(*mpeg1.1*;*mCherry*); TgBAC(*csf1ra*:*GFP*) and *csf1ra^j4e1/j4e1^*; Tg(*mpeg1.1*;*mCherry*); TgBAC(*csf1ra*:*GFP*) fish at multiple timepoints post injury via FACS (Figure 7A-C). This confirmed a significant reduction in the proportion of *mpeg1.1*+ *csf1ra*+ cells in *csf1ra^j4e1/j4e1^*fish at all timepoints in comparison to wildtype fish, except 14 dpi when numbers appeared to recover (Figure 7A; ^69^). Conversely, little difference in the proportion of *mpeg1.1+ csf1ra^dim^* cells was observed at any injury timepoint in either wildtype or *csf1ra^j4e1/j4e1^* fish and a significant difference between the two groups was only observed at 3 dpi (Figure 7B). There was a marked increase in the proportion of *csf1ra*+ cells at all timepoints in *csf1ra^j4e1/j4e1^* fish when compared to wildtype fish and both groups showed a significant increase in the number of *csf1ra*+ cells at 1 dpi when compared to unwounded, which was more marked in *csf1ra^j4e1/j4e1^* fish (Figure 7C).

**Figure 7.**
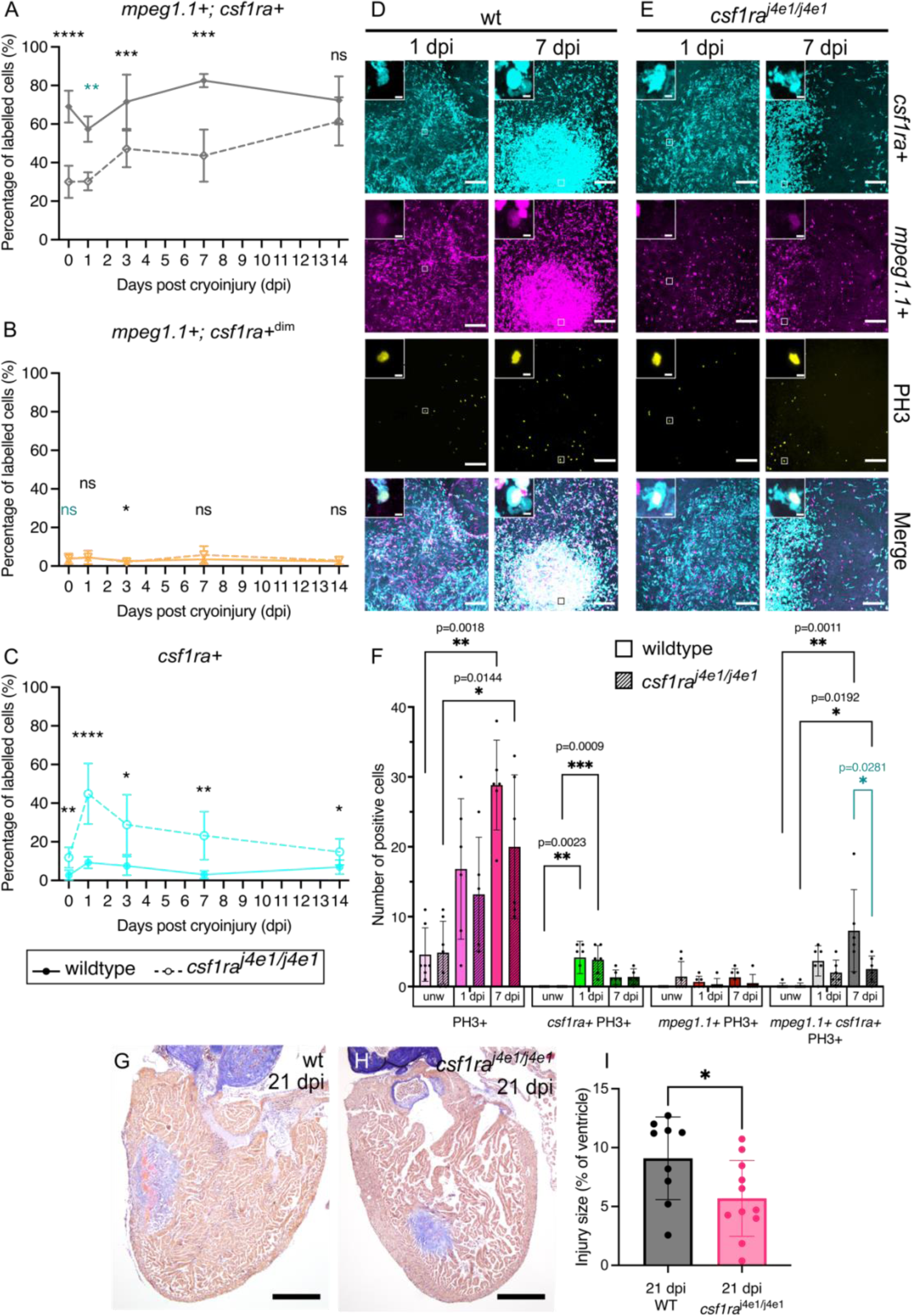
Ventricular *mpeg1.1± csf1ra±* cells proliferate in response to cardiac injury in wildtype and *csf1ra^j4e1/j4e1^*fish. **A-C**) Frequency of each FAC sorted *csf1ra*+ subpopulation expressed as a percentage of labelled cells in wildtype and *csf1ra^j4e1/j4e1^*ventricles at the timepoints indicated. N = 6-8 per timepoint and condition. Statistical analyses = Mann-Whitney pairwise test between experimental groups (blue); Welch’s pairwise test between experimental groups (black). (**D,E**) Representative single channel and merge channel images of wild type (D) and *csf1ra^j4e1/j4e1^*hearts (E) at 1 dpi and 7 dpi following labelling with anti-GFP (*csf1ra*+ - cyan), anti-mCherry (*mpeg1.1*+ - magenta) and anti-PH3 (proliferating cells – yellow) antibodies. Insets show examples of proliferating MNPs identified by the boxes. Scale bars = 100 µm and 5 µm for insets. **F**) Quantification of proliferating (PH3+) cells within each genotype at each timepoint. The total number of PH3+ cells were manually counted, grouped by their co-expression with *mpeg1.1* and *csf1ra*+, per 580 μm x 580 μm x 500 μm (xyz) stack acquired at the border zone. N = 5-7 per condition. Statistical analyses = Mann-Whitney pairwise test between experimental groups (blue); Kruskal-Wallis with Dunn’s multiple comparison tests between all timepoints within each experimental group (black). Comparisons without error bars were not significant. **G-I)** Representative AFOG images of 21 dpi wildtype (G) and *csf1ra^j4e1/j4e1^* (H) hearts showing the composition of the injury site. Quantification of injury area (I) shown as the percentage of the ventricle area. For each fish values were averaged from 3 different slides through the injury. N numbers (wildtype/*csf1ra^j4e1/j4e1^*): 9/11.

To investigate the increase in *csf1ra*+ cells at 1 dpi further, we analysed the rate of proliferation in labelled cells in the heart at 1 and 7 dpi via PH3 immunostaining (Figure 7D-F). As expected, significant numbers of non-MNP PH3+ cells were observed at 7 dpi in both wildtype and *csf1ra^j4e1/j4e1^*fish (Figure 7F). A significant increase in the number of PH3+ *csf1ra*+ cells was observed at 1 dpi in wildtype fish and this was even more marked in *csf1ra^j4e1/j4e1^*fish (Figure 7F). Conversely, very few proliferating *mpeg1.1*+ cells were observed (Figure 7F). Very few PH3+ *mpeg1.1*+ *csf1ra*+ cells were observed at 1 dpi and, although their numbers were significantly increased at 7 dpi in both wildtype and *csf1ra^j4e1/j4e1^* fish, there were significantly less in *csf1ra^j4e1/j4e1^* fish (Figure 7F). Together, this suggests that, as observed in neonatal mice, zebrafish cTMs are capable of expansion via proliferation after injury. And, as the increase in proliferating *csf1ra*+ cells at 1 dpi was higher in *csf1ra^j4e1/j4e1^* fish compared to wildtype this likely contributes to the increased proportion of these cells observed via FACS and further supports the idea that the majority of the remaining myeloid cells in *csf1ra^j4e1/j4e1^* fish are a population of primitive derived cTMs that respond rapidly to injury.

To determine the long-term consequence of the expansion of *csf1ra*+ cells on cardiac regeneration, we analysed the injury area in wildtype and *csf1ra^j4e1/j4e1^*fish at 21 dpi (Figure 7G-I). We have previously reported a reduction in scar deposition at early time points after cardiac injury in *csf1ra^j4e1/j4e1^* fish^24^. Histological analysis at 21 dpi revealed a reduced injury size in *csf1ra^j4e1/j4e1^*fish compared to wildtype controls, although scars were still observed in all fish examined. This suggests that *csf1ra^j4e1/j4e1^* fish retain a smaller injury throughout the repair and regeneration timeline.

## Discussion

Studies in neonatal mice have indicated the importance of cardiac tissue resident macrophages in promoting regeneration after injury^20^. Although previous studies have also revealed crucial roles for macrophage populations in promoting heart regeneration in adult zebrafish^24,29^, we have lacked markers and tools to define the origins and stratify subpopulations of MNPs. Multiple studies have provided important insight into the ontogeny and maintenance of microglia^65,67,70,79^, but much of this hasn’t been investigated in other tissues. Recent studies using single cell sequencing (scRNA-seq) are advancing our understanding of immune cell heterogeneity in the zebrafish heart and other organs^41–43,66,70^ but much of this has been derived from *mpeg1.1* expressing cells and is yet to be translated into new tools to define the distribution of these subpopulations. Our data indicates that using *csf1ra* and *mpeg1.1* transgenic lines it is possible to identify four separate immune cell populations in the homeostatic heart and we have shown that these cells have distinct morphology, distribution, and behaviour in cardiac tissue.

Our data indicate that the embryonic zebrafish heart is seeded with *csf1ra*-expressing macrophages early in development and that the number of these cells remains relatively constant up to juvenile stages, although their distribution alters during that time. Indeed, analysis in *csf1ra^j4e1/j4e1^*and *cmyb*^−/−^ mutants suggests that the early *csf1ra*+ cells are primitive haematopoiesis-derived cells, matching what has been described for beneficial tissue resident populations in mice. Additionally, we observed a population of MNPs that only express *csf1ra* purely on the surface of adult zebrafish hearts and these cells proliferate shortly after injury, again mirroring what is observed in neonatal mice^19^. Collectively, our data suggests these cells are an embryonic-derived population of cTMs that share similarities with those observed in neonatal mice.

In contrast to *csf1ra*+ cTMs, significant numbers of *mpeg1.1*+ macrophages are only observed in the heart from late larval stages onwards. Interestingly, the timing of this *mpeg1.1*+ cell number expansion is the same as that reported in whole larvae^66^ and in microglia^67^ for the onset of significant HSC contribution to circulating macrophages. Indeed, FACS of adult hearts from Tg(*kdrl:cre*); Tg(*actb2:loxP-STOP-loxP-DsR^edexpress^*); Tg(*mpeg1.1:eGFP*) triple transgenic fish, which allow fate-mapping of hemogenic endothelium-derived definitive hematopoietic cells via expression of DsRed^51,67^ suggests that all *mpeg1.1*+ cells are also *DsRed*+ supporting previous findings that suggests all *mpeg1.1* expressing cTMs in the adult heart are definitive-derived^80^ (data not shown). scRNA-seq data suggests differing macrophage phenotypes depending on the tissue of origin with cardiac *mpeg1.1*+ macrophages exhibiting a more pro-reparative phenotype than macrophages in barrier tissues^43^. This further indicates our need to define individual macrophage subpopulations more precisely in order to understand their pro-regenerative roles.

Analysing Tg(*mpeg1.1*;*mCherry*); TgBAC(*csf1ra*:*GFP*) double transgenic fish by FACS revealed an additional double positive population, allowing four separate sub-populations to be identified via these two transgenes. The *mpeg1.1+ csf1ra^dim^* population differed from *mpeg1.1+ csf1ra+* cells in size and could be characterised by loss of *mertka* expression. The absence of the tyrosine kinase receptor MertK, which is involved in phagocytosis^81^, has been described in mammalian classical DCs and in monocytes^18^. DCs have been described in other zebrafish tissues^33^ and only very recently in the heart via scRNA-seq^43^. Interestingly, recent scRNA-seq data suggests large numbers of DCs in the heart^43^ and CNS^70^ of adult zebrafish and that these cells express *mpeg1.1* but have reduced levels of *csf1ra*, supporting our findings. Mammalian DCs are suggested to be smaller than macrophages, mirroring our cytology data, and are most abundant in the atria (specifically the right atrium) rather than the ventricles^82^. Conventional DCs (cDCs) have been shown to expand after MI in mice and depleting the heart of cDCs results in reduced inflammatory activation and improved cardiac function after MI, suggesting these cells may exert a pathological role on the heart following MI^82^. Our data suggests the *mpeg1.1+ csf1ra^dim^* population does not expand after injury to the adult zebrafish heart which may potentially contribute to the regenerative potential of this model.

Our data suggest striking similarities in the formation and maintenance of tissue resident cardiac MNP populations between adult zebrafish and rodents. Our data suggest that *mpeg1.1*+ macrophages are predominant in the adult zebrafish heart and supports previous studies that suggests these cells are all definitive haematopoiesis-derived^80^. We and others have shown that there is considerable expansion of macrophages following injury and this is at least partly due to recruitment from cells outside the heart^24,29^. Although the *csf1ra*+ cTMs proliferate after injury this is not sufficient to account for the large expansion in cardiac macrophages observed, further suggesting that significant numbers of monocyte-derived macrophages are recruited, as observed in adult mammals. Indeed, reduced definitive/monocyte-derived macrophage infiltration in *csf1ra^j4e1/j4e1^*fish early after injury (3 dpi) could explain our previous observations that these mutants exhibit reduced macrophage pro-inflammatory activation and reduced scar deposition^24^. However, at later stages of regeneration *csf1ra^j4e1/j4e1^* fish do exhibit cardiac scarring, albeit to a lesser extent than wildtype fish. This could suggest that the monocyte-derived macrophages that infiltrate the injury also contribute to regeneration, unlike in mammals, and future work should concentrate on expanding our understanding of the potentially pro-regenerative functions of these cells.

## Funding

This work was supported by the British Heart Foundation (PG/21/10646; FS/15/2/31225 to R.J.R), a Wellcome Trust–funded Ph.D. studentship (108907/Z/15/Z to B.R.M) and the Fonds de la Recherche Scientifique (FNRS) (J.0202.22 to V.W). We acknowledge funding for bioimaging equipment from a BBSRC Alert 13 capital grant (BB/L014181/1), BBSRC Alert 19 equipment grant (BB/T017597/1), MRC funding of a pre-clinical In-vivo functional imaging platform for translational regenerative medicine, and an MRC grant (MRC MC_PC_MR/X01391X/1).

## Acknowledgements

We wish to thank Professor Stephen Renshaw and Catherine Loynes at the University of Sheffield for assisting with the TgBAC(*csf1ra:Gal4i);* Tg(*UAS:kaede*) experiment. We thank the Wolfson Bioimaging Facility at the University of Bristol, and particularly Stephen Cross and Dominic Alibhai for imaging expertise, and the University of Bristol Faculty of Biomedical Sciences Flow Cytometry Facility for cell sorting. We also thank Marianne Caron at the Université Libre de Bruxelles (ULB) for technical assistance.

## Notes

### Competing Interest Statement

The authors have declared no competing interest.

