## Supplementary material for "Defining mononuclear phagocyte distribution and behaviour in the zebrafish heart"

Supplementary Table 1

| **A) Summary of human and zebrafish orthologs to commonly used murine MNP marker** | | | | |
|---|---|---|---|---|
| **Mouse** | | | **Orthology** | |
| **Gene** | **Expression** | **Function** | **Human** | **Zebrafish** |
| *Csf1r* | - Expressed by all cells of the **MNP** system | Cell surface receptor for M-CSF/CSF-1 which is required for the survival, proliferation and differentiation of MNPs | *CSF1R* (11/11) | *csf1ra* (10/11)<br>*csf1rb* (2/11) |
| *Csf2r* | - Expressed by all cells of the **MNP** system<br>- **Macrophages,** T-cells, B-cells, endothelial cells, vascular smooth muscle cells neutrophils, eosinophils, and monocytes | Plays major role as a pro-inflammatory cytokine and for positive regulation of leukocyte proliferation. Haematologic cell growth factor, stimulating blood stem cells to produce granulocytes and monocytes. | *CSF2RB* (11/11) | *csf2rb* (7/11) |
| *Csf3r* | - Expressed by neutrophils, **monocytes and macrophages**, HSPCs, and a subset of B lymphocytes and NK cells | Essential for the development and maturation of granulocytic neutrophils from hematopoietic precursors in the bone marrow. | *CSF3R* (11/11) | *csf3r* (10/11) |
| *Ccr2* | - High expression by **classical monocytes**<br>- Low expression by non-classical monocytes and yolk-sac derived/embryonic, tissue resident (including cardiac) macrophages. | Required for monocyte mobilisation and recruitment to tissues expressing its ligand, Ccl2. | *CCR2* (5/11) | *ccr2* (5/11) |
| *Cx3cr1* | - Highly expressed by **tissue-resident macrophages**<br>- High expression on **non-classical monocytes**; low expression by classical monocytes | Chemokine receptor | *CX3CR1* (10/11) | *ccr2* (1/11) |

| | | | | |
|---|---|---|---|---|
| *Ly6C1* | - High expression by **classical monocytes** which differentiate into monocyte derived macrophages.<br>- Low expression by non-classical monocytes | Largely unknown. Reported to have acetylcholine receptor binding activity. | *LY6H (*3/11) | - |
| *Adgre1 (F4/80)* | - **Monocytes and macrophages**<br>- High expression on **tissue resident macrophages** | Cell adhesion molecule | *ADGRE1* (11/11) | *adgre10* (3/10)<br>*adgre5b.2* (3/10) |
| *Itgam (Cd11b)* | - Expressed on all **MNPs** in addition to granulocytes and natural killer cells | Integrin, cell adhesion molecule | *ITGAM* (9/11) | *itgam.1* (8/11)<br>*itgam.2* (1/11) |
| *Cd68* | - **Macrophages** | Scavenger receptor | *CD68* (9/11) | *cd68* (2/11) |
| *Sell (Cd62l)* | - Expressed by **classical monocytes** | L-selectin, cell adhesion molecule | *SELL* (11/11) | - |
| *Itgax (Cd11c)* | - **Conventional DCs** | Integrin, cell adhesion molecule | *ITGAX* (11/11) | *itgam.1* (10/11) |
| *Mrc1 (Cd206)* | - Highly expressed by **classically activated/M1 macrophages**<br>- Low expression on alternatively activated/M2 macrophages | Pattern recognition receptor | *MRC1* (11/11) | *mrc1a* (10/11)<br>*mrc1b* (4/11) |
| *Mertk* | - Highly expressed by **macrophages** | Involved in phagocytosis | *MERTK* (10/11) | *mertka* (10/11) |
| *Timd4* | - Expressed by populations of **resident cardiac macrophages** | Involved in apoptotic cell clearance | *TIMD4* (11/11) | *timd4* (2/11) |

**B) Summary of human and murine orthologs to commonly used zebrafish MNP markers**

| Zebrafish | | | Orthology | |
|---|---|---|---|---|
| **Gene** | **Expression** | **Function** | **Human** | **Mouse** |
| *csf1ra* | As shown in **Error! Reference source not found.** | | | |
| *mpeg1.1* | - Macrophages, unknown in other MNP populations | Pore-forming perforin involved in pathogen responses | *MPEG1* (2/11) | *Mpeg1* (2/11) |
| *mpeg1.2* | - MNPs | Pore-forming perforin involved in pathogen responses | *MPEG1* (10/11) | *Mpeg1* (10/11) |
| *mfap4* | - Expressed in monocytes and macrophages, unknown expression in DCs | Predicted antigen binding activity | *MFAP4* (8/11) | *Mfap4* (8/11) |
| *marco* | - Expressed in macrophages | Scavenger receptor | *MARCO* (7/11) | *Marco* (7/11) |
| *mhc2dab* | - Highly expressed by MNPs | Antigen presentation molecule | Functional orthologs are annotated | |

| | | | | |
|---|---|---|---|---|
| *apoeb* | - Highly expressed by microglia | Involved in lipid homeostasis | *APOE* (10/11) | *Apoe* (11/11) |
| *acod1 (irg1)* | - Activated macrophages | Bactericidal roles | *ACOD1* (10/11) | *Acod1* (10/11) |
| *mertka* | - Phagocytic cells | Involved in signalling pathways required for phagocytosis | *MERTK* (9/11) | *Mertk* (10/11) |

**Supplementary Table 1. Summary of human, murine and zebrafish MNP marker orthologs. A)** Classical murine marker genes of MNP populations were compiled from a range of published literature (Epelman et al., 2014, Dick et al., 2019, Nahrendorf et al., 2007, Shiraishi et al., 2016, Reynolds and Haniffa, 2015) and humans and zebrafish orthologs investigated using The Alliance database (https://www.alliancegenome.org/). Numbers in brackets indicate the total number of databases searched and the number that gene as an ortholog. Dashes indicate where no ortholog was identified. **B)** Classical zebrafish marker genes of MNP populations were compiled from a range of published literature (Zakrzewska et al., 2010, Benard et al., 2015, Sanderson et al., 2015, Walton et al., 2015, Peri and Nüsslein-Volhard, 2008, Wittamer et al., 2011) and the reverse analysis was applied as above.

**Supplemental Video 1.** 3D Imaris render of individual *mpeg1.1*+ (magenta) cells on the surface of the ventricle associated with *myl7*+ (grey) cardiomyocytes in an unwounded Tg(*mpeg1.1:GFP*); Tg(*myl7:HRAS-mCherry*) heart. Scale bar = 5 – 20 µm as indicated per frame.

**Supplemental Video 2.** 3D Imaris render of individual *mpeg1.1*+ (magenta) cells on the surface of the ventricle associated with *kdrl*+ (grey) endothelial cells in an unwounded *Tg(mpeg1.1:GFP); Tg(kdrl:mCherry-CAAX)* heart. Scale bar = 5 – 10 µm as indicated per frame.

**Supplemental Video 3.** 3D projection of a cleared, unwounded Tg*(mpeg1.1:mCherry);* TgBAC(*csf1ra:GFP)* ventricle. *mpeg1.1* is shown in magenta; *csf1ra* is shown in cyan. Scale bar = 100 µm.

**Supplemental Video 4.** 3D projection of a cleared, unwounded *csf1ra*$^{j4e1/j4e1}$; Tg*(mpeg1.1:mCherry);* TgBAC(*csf1ra:GFP)* mutant ventricle. *mpeg1.1* is shown in magenta; *csf1ra* is shown in cyan. Scale bar = 100 µm.

**Supplemental Video 5.** *Ex vivo* imaging of an unwounded Tg(*mpeg1.1:mCherry*); TgBAC(*csf1ra:GFP*) ventricle using spinning disk confocal microscopy. The video shows maximum projections of *csf1ra:GFP* (cyan), *mpeg1.1:mCherry* (magenta), and merged channel images. Frames are at 2.5 minute intervals for 100 minutes. Scale bar = 20 µm.
